# Critical role of Gα12 and Gα13 proteins in TGF-β-induced myofibroblast differentiation

**DOI:** 10.1101/2024.05.29.596473

**Authors:** Eleanor B. Reed, Albert Sitikov, Robert B. Hamanaka, Rengül Cetin-Atalay, Gökhan M. Mutlu, Alexander A. Mongin, Nickolai O. Dulin

**Author notes:** To whom correspondence should be addressed: Nickolai O. Dulin, The University of Chicago Department of Medicine, Section of Pulmonary and Critical Care Medicine, 5841 S. Maryland Ave, MC6076, Chicago, IL 60637.

## Abstract

Myofibroblast differentiation, characterized by accumulation of cytoskeletal and extracellular matrix proteins by fibroblasts, is a key process in wound healing and pathogenesis of tissue fibrosis. Transforming growth factor-β (TGF-β) is the most powerful known driver of myofibroblast differentiation. TGF-β signals through transmembrane receptor serine/threonine kinases that phosphorylate Smad transcription factors (Smad2/3) leading to activation of transcription of target genes. Heterotrimeric G proteins mediate a distinct signaling from seven-transmembrane G protein coupled receptors, not commonly linked to Smad activation. We asked if G protein signaling plays any role in TGF-β-induced myofibroblast differentiation, using primary cultured human lung fibroblasts. Activation of Gαs by cholera toxin blocked TGF-β-induced myofibroblast differentiation without affecting Smad2/3 phosphorylation. Inhibition of Gαi by pertussis toxin, or siRNA-mediated combined knockdown of Gαq and Gα11 had no significant effect on TGF-β-induced myofibroblast differentiation. A combined knockdown of Gα12 and Gα13 resulted in a drastic inhibition of TGF-β-stimulated expression of myofibroblast marker proteins (collagen-1, fibronectin, smooth-muscle α-actin), with siGα12 being significantly more potent than siGα13. Mechanistically, a combined knockdown of Gα12 and Gα13 resulted in a substantially reduced phosphorylation of Smad2 and Smad3 in response to TGF-β, which was accompanied by a significant decrease in the expression of TGFβ receptors (TGFBR1, TGFBR2) and of Smad3 under siGα12/13 conditions. In conclusion, our study uncovers a novel role of Gα12/13 proteins in the control of TGF-β signaling and myofibroblast differentiation.

## Introduction

Transforming growth factor-β (TGF-β) is a pleotropic cytokine with multiple cell-specific functions. TGF-β was originally called “transforming” because it enhanced anchorage-independent growth of normal rat kidney (NRK) cells on soft agar (the commonly used assay for cell transformation) in response to TGF-α or epidermal growth factor [1, 2]. Subsequently it was found that TGF-β inhibited anchorage-dependent growth of NRK cells and of multiple human tumor cell lines; and it has been recognized as an inhibitor of cell cycle progression and cell proliferation [3]. Through numerous studies, it is now established that TGF-β controls fundamental cellular processes such as cell proliferation, survival, hypertrophy, senescence, epithelial-to-mesenchymal transition, cell differentiation; and it is implicated in a variety physiological and pathological processes [4]. This study focuses on signaling mechanisms that mediate one of the functions of TGF-β – differentiation of fibroblasts to myofibroblasts.

Myofibroblasts are modified fibroblasts, originally characterized by the presence of a well-developed contractile apparatus, and the formation of robust actin stress fibers containing smooth muscle α-actin (SMA) isoform normally expressed in smooth muscle cells [5, 6]. Myofibroblast produce extracellular matrix proteins such as fibronectin, multiple isoforms of collagen and other proteins implicated in matrix remodeling [7-9]. Over many decades of research, myofibroblasts have been recognized as the key cells in wound healing and pathogenesis of tissue fibrosis [10, 11].

TGF-β is the most powerful known driver of myofibroblast differentiation [12]. TGF-β signals through transmembrane-receptor serine/threonine kinases that phosphorylate Smad transcription factors (Smad2/3), leading to their heteromerization with a common mediator Smad4, nuclear translocation of Smad2/3/4 complex and activation of transcription of target genes [13, 14]. G protein coupled receptors (GPCRs), the largest receptor family regulating the function of all mammalian cells, transduce extracellular signals through heterotrimeric G proteins, with Gα and Gβγ subunits controlling the activity of specific target proteins [15]. Four functionally distinct types of Gα subunits have been identified: Gαs, Gαi, Gαq/11, and Gα12/13 [16]. Gαs activates adenylyl cyclase to produce cAMP, whereas Gαi inhibits this enzyme [17].

Gαq/11 activate phospholipase Cβ [18, 19] generating two second messengers – inositol trisphosphate and diacylglycerol that stimulate calcium release from endoplasmic reticulum and activate protein kinase C, respectively [20]. In addition, Gβγ proteins can also activate PLCβ isoforms [21]. Gα12/13 stimulate Rho family of small GTPases through a direct recruitment of specific guanine exchange factors for RhoA small GTPase [22-25].

Litte is known about the crosstalk between TGF-β and G protein signaling in the context of myofibroblast differentiation. Agonists coupled to Gαs (i.e. prostaglandin E2, prostacyclin, adrenomedullin) have been shown by us and others to inhibit TGF-β-induced myofibroblast differentiation through a protein kinase A (PKA)-dependent mechanism [26-29]; however, the role of Gαs has not been evaluated in these studies. Better understanding of the role of Gαs is important, given that PKA can be stimulated through other mechanisms, including the Gβγ-mediated one [30]. GPCR agonists acting through Gαi, Gαq/11, and Gα12/13 (i.e. lysophosphatidic acid, sphyngosine-1-phosphate) have been reported to promote myofibroblast differentiation [31, 32]; however, the role of specific G proteins was not identified. A link between TGF-β and G protein signaling has been reported, wherein GPCR ligands (angiotensin II, thrombin, etc.) promote TGF-β synthesis [33, 34] or a release of active TGF-β from the pericellular matrix [35]. However, a direct role of G proteins in TGF-β signaling in the context of myofibroblast differentiation has not been investigated. In this study, we sought to determine the role that G proteins play in TGF-β signaling in the context of myofibroblast differentiation.

## Materials and Methods

### Primary culture of human lung fibroblasts (HLFs)

HLFs were isolated from the lungs of patients with IPF shortly after their removal during lung transplantation at the University of Chicago under the IRB protocol #14514A. Human lung tissue samples were placed in DMEM with antibiotics. Lung tissue was minced to ∼1 mm^3^ pieces, washed, and plated on 10-cm plates in growth media containing DMEM supplemented with 10% FBS and antibiotics. The media was changed twice a week. After approximately 2 weeks, the explanted and amplified fibroblasts were trypsinized, cleared from tissue pieces by sedimentation, and further amplified as passage

1. Unless indicated, cells were grown in a growth media for 24 hours, starved in DMEM containing 0.1% FBS for 48 hours, and treated with desired drugs for various times as indicated in the figure legends. Primary cultures were used from passage 3 to 8.

### siRNA transfection

HLFs were plated at a density of 0.4 x10^5^ cells per well (24-well plates) and were grown for 24 hours. Cells were then transfected with total of 10 nM desired siRNA using Lipofectamine RNAiMAX Reagent (ThermoFisher Scientific, Waltham, MA) according to the standard protocol, and kept in growth media for additional 24 hours, followed by serum starvation in 0.1% FBS for 48 hours, and then treatment with TGF-β for desired times.

### Western blotting

Cells were lysed in a buffer containing 8 M deionized urea, 1% SDS, 10% glycerol, 60mM Tris-HCl, pH 6.8, 0.02% pyronin Y, and 5% β-mercaptoethanol. Lysates were sonicated for 5 seconds. Samples were then subjected to polyacrylamide gel electrophoresis and Western blotting with desired primary antibodies and corresponding horseradish peroxidase (HRP)-conjugated secondary antibodies and developed by chemiluminescence reaction. Digital chemiluminescent images below the saturation level were obtained with a LAS-4000 analyzer, and the light intensity was quantified using Multi Gauge software (Fujifilm, Valhalla, NY).

Primary antibodies were validated by molecular weight of target proteins and by siRNA-mediated knockdown.

### Materials

Recombinant TGF-β (T7039), Cholera toxin (227036) and Pertussis toxin (516560) were from Millipore-Sigma. The following antibodies for Western blotting were from Millipore-Sigma: smooth muscle α-actin (A5228, 10,000X), β-actin (A5441, 10,000X), α-tubulin (T6074, 10,000X). Fibronectin antibody (610077, 1,000X) was from BD Transduction. Antibodies against human collagen-1A1 (sc-28657, 1,000X), Gα12 (sc-409, 200X), TGFBR2 (sc-400) were from Santa Cruz Biotechnology. Antibodies against Smad2 (L1603, 1,000X), phospho-Smad2-Ser465/467 (138D4, 1,000X), phospho-Smad3-Ser423/425, 1,000X) were from Cell Signaling Technology. Gα13 antibody (GTX32613, 1,000X) was from GeneTex. Smad3 antibody (06-920, 1,000X) was from Upstate Biotechnology. TGFBR1 antibody (AB235578, 1,000X) was from Abcam. Secondary HRP-conjugated antibodies for western blotting (1:3,000 dilution) were from Millipore-Sigma (40-139-32 - anti-rabbit IgG, 40-125-32 - anti-mouse IgG).

### Statistical Analysis

In this study, a replicate (n) represents an independently plated and treated HLF culture. All individual data points are presented in figures along with mean values ± standard deviation (SD). Results were analyzed for normal distribution using a Shapiro-Wilk test. Normally distributed data were further statistically compared using one-way ANOVA with the Tukey honest significant difference post hoc correction for multiple comparisons. Values of p<0.05 were considered statistically significant. All statistical analyses were performed in Prism v. 10.2.3 (GraphPad Software, Boston, MA),

## Results

### Activation of Gαs by cholera toxin blocks TGF-β-induced myofibroblast differentiation without affecting Smad2/3 phosphorylation

Previous studies by us and others demonstrated regulation of TGF-β-induced myofibroblast differentiation by agonists acting in part through Gαs-coupled GPCRs [26-29]; however, a direct role of Gαs has not been carefully investigated.

Cholera toxin (CTX), which ADP-ribosylates and inhibits GTPase activity of Gαs, is recognized as a powerful and highly specific activator of Gαs [36]. Therefore, we used CTX as a tool to investigate the effect of Gαs activation on TGF-β-induced myofibroblast differentiation. As shown in Fig. 1A, CTX, applied shortly following TGF-β treatment for 48 hours, abolished accumulation of myofibroblast marker proteins, collagen 1A1 (Col1A1), fibronectin (FN) and smooth muscle α-actin (SMA). Pretreatment of cells with CTX for 2 hours did not affect Smad2/3 phosphorylation induced by acute (30 minute) TGF-β treatment (Fig. 1B). Stimulation of Gαs by CTX was confirmed by western blotting with “protein kinase A (PKA) substrate” (PKAS) antibodies that recognize proteins phosphorylated by PKA - a downstream effector of Gαs. Specificity of PKAS antibodies was previously demonstrated by the expression of a specific PKA inhibitor protein, PKI [37]. These data indicate that active Gαs abolishes TGF-β-induced myofibroblast differentiation without affecting proximal TGF-β signaling.

**Figure 1.**
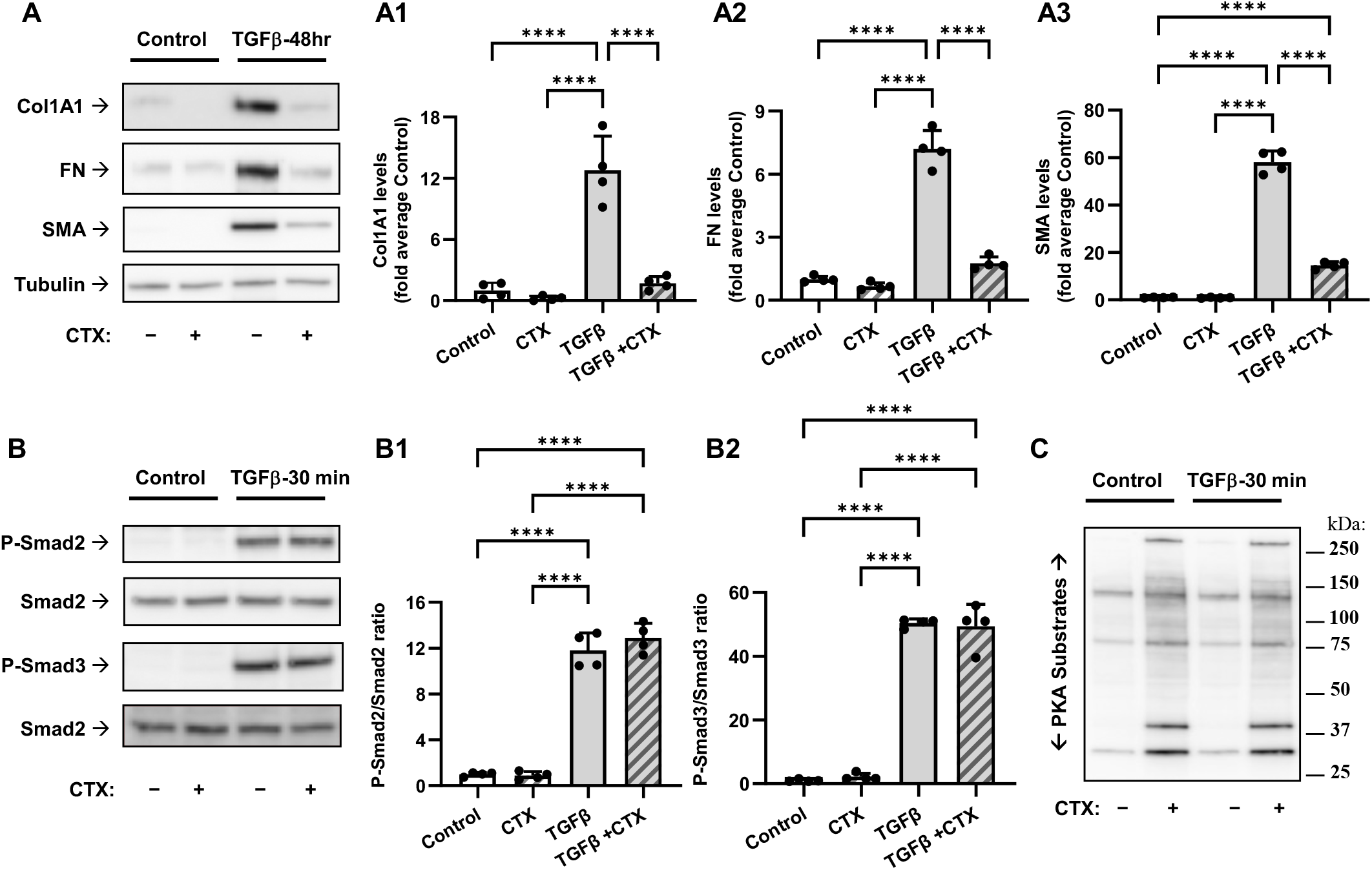
Activation of Gas by cholera toxin blocks TGF-b-induced myofibroblast differentiation without affecting Smad2/3 phosphorylation. **A**, Representative images and quantification of western blot analyses of myofibroblast markers in human lung fibroblasts (HLF) treated with/without 1 ng/ml TGF-b and/or 1 mg/ml cholera toxin (CTX) for 48 h. Cell lysates were analyzed using antibodies recognizing collagen 1A1 (Col1A1, **A1**), fibronectin (FN, **A2**), and smooth muscle actin (SMA, **A3**). The relative immunoreactivity values were normalized to the ubulin immunoreactivity in the same protein lysate (not shown) and then to the average immunoreactivity or control samples. Data are the mean values ± SD from 4 independent cultures per treatment. ****p<0.001, one-way ANOVA with Tukey correction for multiple comparisons. **B**, Representative images and quantification of western blot analyses of Smad2/3 phosphorylation in HLF pretreated with or without 1 mg/ml CTX following 30 – min exposure to ng/ml TGF-b. Cell lysates were analyzed using antibodies recognizing Smad2/3 and their phosphorylated forms. pSmad/Smad ration was determined in each sample for Smad2 (B**1**) and Smad3 (**B2**), and smooth muscle actin (SMA, **A3**). Data are the mean values ± SD from 4 independent cultures per treatment. ****p<0.001, one-way ANOVA with Tukey correction for multiple comparisons. **C**, HLF were pretreated with 1 mg/ml CTX for 2 hours, followed by treatment with TGFb for 30 minutes. Cell lysates were analyzed by western blotting with desired antibodies. **C**, Representative western blot image of HLF treated as in *B* and then probed with antibody recognizing phosphorylated PKA substrates.

### Inhibition of Gαi by pertussis toxin does not affect TGF-β-induced myofibroblast differentiation

We then focused on the role of Gαi, using pertussis toxin (PTX), which ADP-ribosylates and blocks the activity of Gαi through inhibition of GDP to GTP exchange by Gαi [38, 39]. Pretreatment of HLFs with PTX had no significant effect on TGF-β-induced expression of Col1A1, FN and SMA (Fig. 2A). We have previously established that Gαi-coupled Gβγ mediates phosphorylation of extracellular signal regulated kinases ERK1/2 by endothelin-1 (ET-1) in vascular smooth muscle cells [30]. Therefore, we confirmed that PTX was effective in the inhibition of Gαi in HLFs by demonstration that PTX pretreatment abolished ET1-induced phosphorylation of ERK1/2 (Fig. 2B). Together, these data suggest that Gαi may not be involved in TGF-β-induced myofibroblast differentiation.

**Figure 2.**
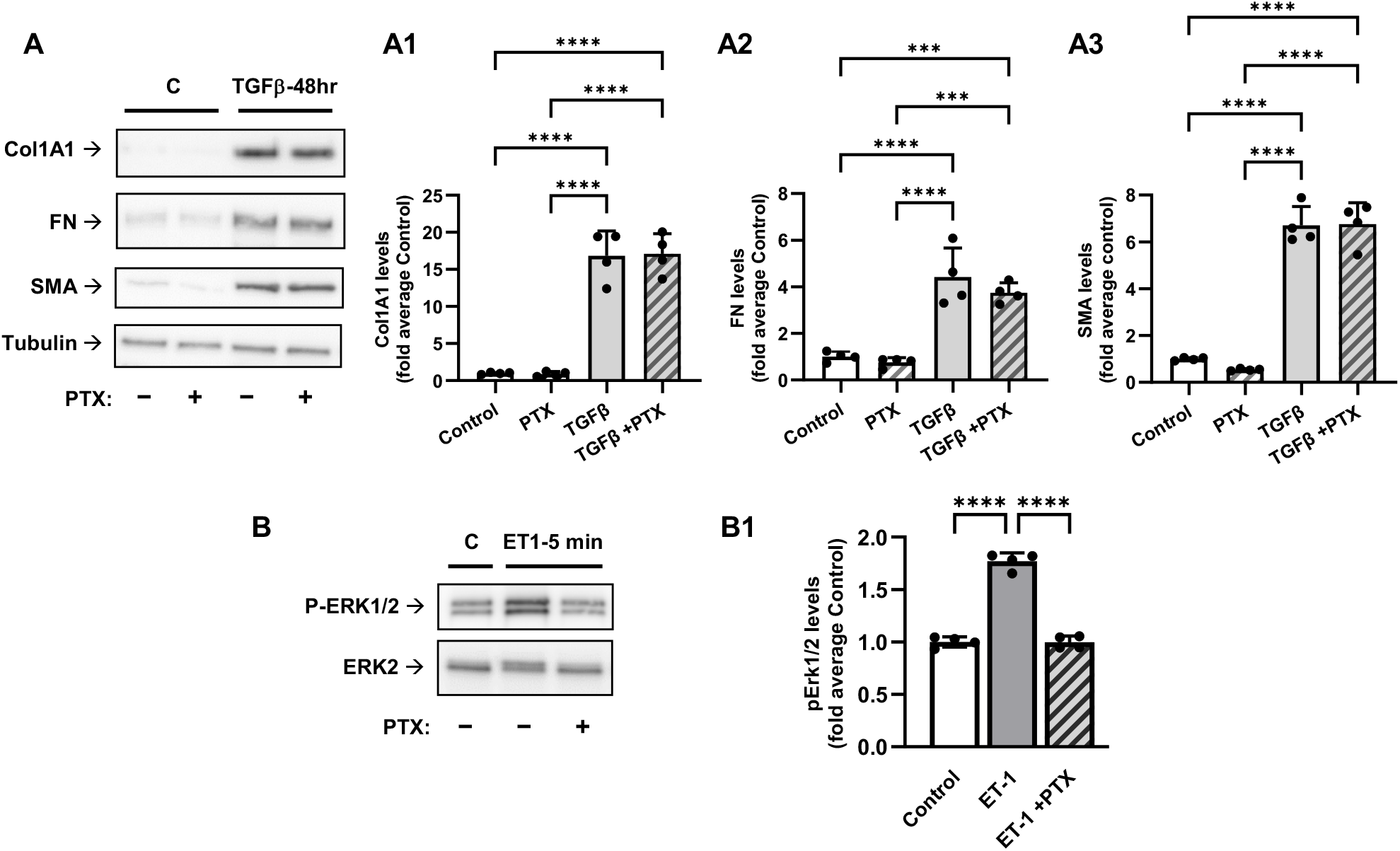
Inhibition of Gαi by pertussis toxin does not affect TGFβ-induced myofibroblast differentiation. **A**, Representative images and quantification of western blot analyses of HLF pretreated with 100 ng/ml pertussis toxin (PTX) overnight, followed by treatment with 1 ng/ml TGFβ for 48 hours. Cell lysates were analyzed using antibodies recognizing collagen 1A1 (Col1A1, **A1**), fibronectin (FN, **A2**), and smooth muscle actin (SMA, **A3**). The relative immunoreactivity values were normalized to the tubulin immunoreactivity in the same protein lysate (not shown) and then to the average immunoreactivity or control samples. Data are the mean values ± SD from 4 independent cultures per treatment. ***p<0.001; ****p<0.001, one-way ANOVA with Tukey correction for multiple comparisons. **B**, Representative images of western blot analyses of the PTX pretreated HFL with or without subsequent 5-min treatment with 100 nM endothelin-1 (ET1). Cell lysates were probed with antibodies recognizing pErk1/2 or total Erk2. **B1**, Quantification of experiments presented in B. ****p<0.001, one-way ANOVA with Tukey correction for multiple comparisons.

### Knockdown of Gαq/11 does not affect TGF-β-induced myofibroblast differentiation

To assess the role of Gαq/11, we used the siRNA approach. A combined knockdown of ubiquitous Gαq and Gα11 with corresponding siRNAs resulted in a 70% decrease in the expression of each protein in the presence or absence of TGF-β, as assessed by western blotting with antibodies that recognize both Gαq and Gα11 (Fig. 3). Under the same treatment conditions, Gαq/11 knockdown had no significant effect on TGF-β-induced myofibroblast differentiation, suggesting a possible lack of the role of Gαq/11 in this process.

**Figure 3.**
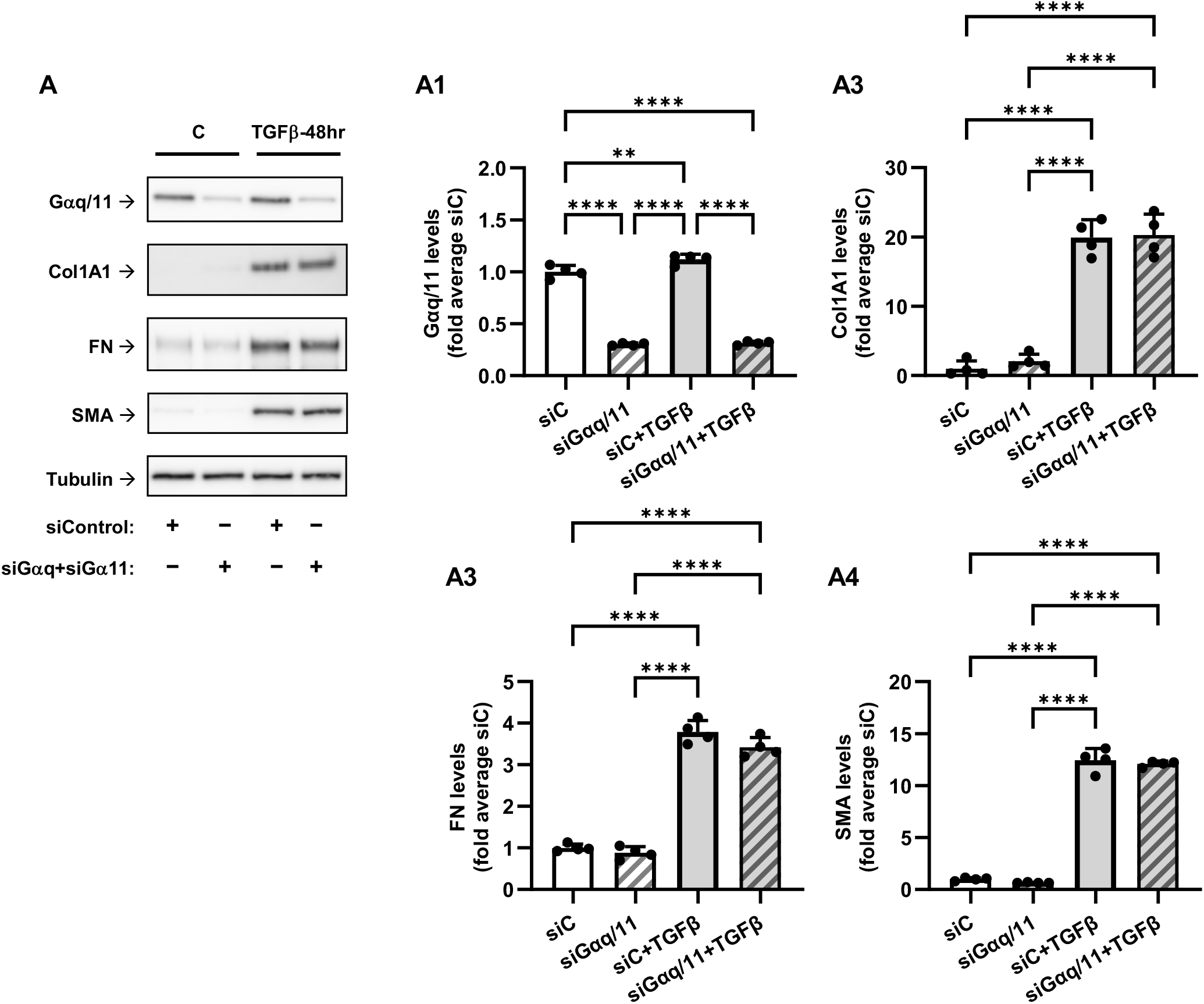
Knockdown of Gαq and Gα11 does not affect TGFβ-induced myofibroblast differentiation. **A**, Representative images and quantification of western blot analyses of HLF transfected with control siRNA (siC) or with siRNAs targeting Gαq and Gα11, starved for 48 h, followed by the treatment with 1 ng/ml TGFβ for 48 hours. Cell lysates were analyzed using antibodies recognizing Gαq/Gα11 (**A1**), collagen 1A1 (Col1A1, **A2**), fibronectin (FN, **A3**), and smooth muscle actin (SMA, **A4**). The relative immunoreactivity values were normalized to the tubulin immunoreactivity in the same protein lysate (not shown) and then to the average immunoreactivity or siC-treated control samples. Data are the mean values ± SD from 4 independent cultures per treatment. **p<0.01; ****p<0.001, one-way ANOVA with Tukey correction for multiple comparisons.

### Knockdown of Gα12 and Gα13 attenuates the TGF-β-induced myofibroblast differentiation in a synergistic fashion

We then examined the role of Gα12 and Gα13 in TGF-β-induced myofibroblast differentiation also using siRNA approach. As shown in Figure 4, knockdown of Gα12 and of Gα13 achieved up to 85% and 80% reduction in corresponding protein expression levels in HLFs. Knockdown of Gα12 resulted in a moderate inhibition of TGF-β-induced expression of Col1A1, FN and SMA by 30%, 25%, and 10%, respectively. Knockdown of Gα13 had no significant effect on the expression of these proteins. However, a combined knockdown of Gα12 and Gα13 significantly decreased TGF-β-induced increases in protein levels of Col1A1, FN and SMA by 80%, 100%, and 60%, respectively (Fig. 4). These data suggest that both Gα12 and Gα13 proteins are required for TGF-β-induced myofibroblast differentiation, with Gα12 being potentially of higher importance.

**Figure 4.**
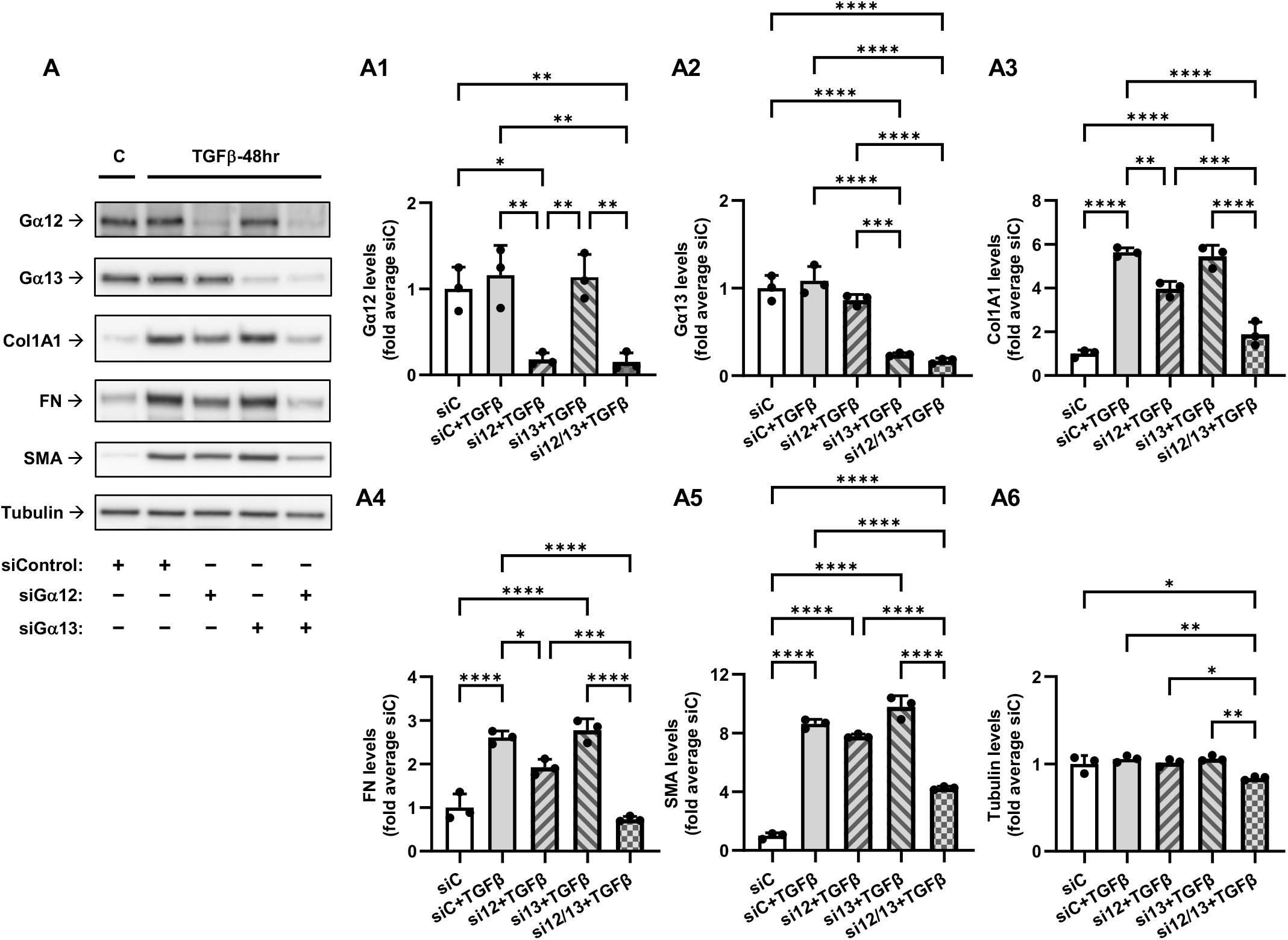
Knockdown of Gα12 and Gα13 attenuates the TGFβ-induced myofibroblast differentiation in a synergistic fashion. **A**, Representative images and quantification of western blot analyses of HLF transfected with control siRNA or with siRNAs for Gα12, Gα13, or with combination of both, overnight. Cells were subsequently serum starved for 48 hours and further treated with 1 ng/ml TGFβ for 48 hours. Cell lysates were analyzed using antibodies recognizing Ga12 (**A1**), Ga13 (**A2**), collagen 1A1 (Col1A1, **A3**), fibronectin (FN, **A4**), smooth muscle actin (SMA, **A5**), or housekeeping tubulin **(A6**). The relative immunoreactivity values were normalized to the tubulin immunoreactivity in the same protein lysates and then to the average immunoreactivity or siNC samples. Data are the mean values ± SD from 3 independent cultures per treatment. *p<0.05, **p<0.01; ***p<0.001; ****p<0.001, one-way ANOVA with Tukey correction for multiple comparisons.

### Combined knockdown of Gα12 and Gα13 inhibits TGF-β-induced Smad2/3 phosphorylation

To begin understanding the mechanism for the role of Gα12/13 in a control of TGF-β-induced myofibroblast differentiation, we tested the effect of a combined Gα12/13 knockdown on TGF-β-induced phosphorylation of Smad2 and Smad3 - an initial event in TGF-β receptor signaling. As shown in Figure 5, Gα12/13 knockdown clearly inhibited TGF-β-induced phosphorylation of both Smad2 and Smad3, as assessed by Western blotting with corresponding phospho-Smad antibodies. The total levels of Smad2 were moderately but significantly decreased by 20%, whereas Smad3 expression was significantly reduced by up to 40% under Gα12/13 knockdown conditions. Despite a decrease in total expression levels of Smad2/3, normalized data indicated a substantial and highly significant reduction of P-Smad2/Smad2 (70%) and of P-Smad3/Smad3 (55%) ratios under siGα12/13 treatment. These data suggested that a decrease in Smad2/3 phosphorylation could not be solely explained by a reduction of Smad2/3 levels. Therefore, we examined TGF-β receptor levels and observed a highly significant reduction in the expression of TGFBR1 (43%) and to a lesser extent of TGFBR2 (23%) in HLFs treated with siGα12/13 compared with HLFs treated with control siRNA (Fig. 5). Together, these data suggest that Gα12/13 are required for TGF-β-induced Smad2/3 phosphorylation and they control the expression of TGF-β receptors and Smad2/3 proteins.

**Figure 5.**
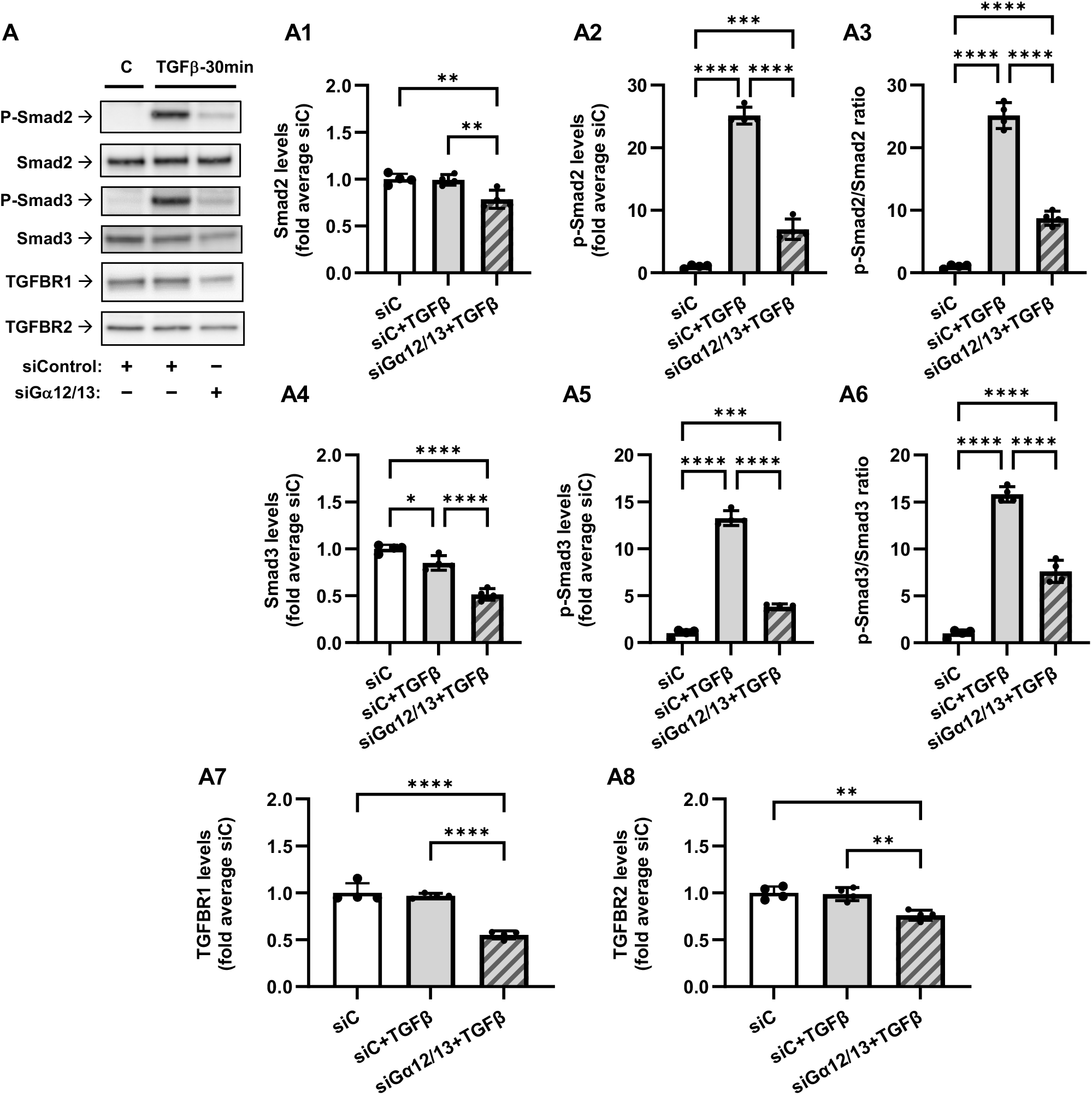
Combined knockdown of Gα12 and Gα13 inhibits TGFβ-induced Smad2/3 phosphorylation and reduces TGFB receptor levels. **A**, Representative images and quantification of western blot analyses of transfected with control siRNA (siC) or combination of siRNAs targeting Gα12 and Gα13 overnight. Cells were then serum starved for 48 hours and additionally treated with 1 ng/ml TGFβ for 30 min. Cell lysates were analyzed using antibodies recognizing p-Smad2, Smad2, p-Smad3, Smad3, TGFBR1, or TGFBR2. Quantifications show immunofluorescence levels of Smad2 (**A1**), p-Smad2 (**A2**), ratios of p-Smad2/Smad2 (**A3**), levels of Smad3 (**A4**), p-Smad3 (**A5**), ratios of p-Smad3/Smad3 (**A3**), and relative expression levels of TGFBR1 (**A7**), and TGFBR2 (**A8**). The relative immunoreactivity values were normalized to the tubulin immunoreactivity in the same protein lysates (not shown) and then to the average immunoreactivity in the siC samples. Data are the mean values ± SD from 4 independent cultures per treatment. *p<0.05; **p<0.01; ***p<0.001; ****p<0.001, one-way ANOVA with Tukey correction for multiple comparisons.

## Discussion

The major finding of this study is the discovery that Gα12/13 proteins are required for TGF-β-induced myofibroblast differentiation, and they control proximal TGF-β signaling (Smad2/3 phosphorylation) (Figs. 4, 5). The important question is whether TGF-β can indirectly activate G protein signaling, specifically that of Gα12/13. We and others have previously demonstrated that TGF-β, through Smad-dependent gene transcription, recruits RhoA signaling, actin polymerization and activation of a transcription factor, serum response factor (SRF), for the induction of SMA expression in fibroblasts [28, 41, 42]. Given that RhoA pathway is activated by Gα12/13 [22-25], it is reasonable to propose that Gα12/13 are activated at some point through TGF-β signaling. The question is which GPCRs may be activated during TGF-β signaling. Some candidate GPCRs are worthy of consideration based on current reports. It was shown that lysophosphatidic acid (LPA), sphingosine-1-phospphate (S1P) and thrombin cooperate in human dermal fibroblasts with TGF-β to induce extracellular matrix synthesis, myofibroblast marker expression and cytokine secretion [43]. Sphingosine-1 phosphate (S1P) receptor signaling was shown to be important for TGF-β-induced myofibroblast differentiation in a number of studies [44-46]. Last but not least, TGF-β induces endothelin-1 (ET1) expression [47], although it may also downregulate ET1 receptors [48]. More than 30 GPCRs have been reported to couple to Gα12/13 [49]; hence identification of critical GPCRs mediating TGF-β signaling related to myofibroblast differentiation needs further investigation and is of potential therapeutic importance for treatment of tissue fibrosis.

We also observed that while a combined knockdown of Gα12/13 abolished TGF-β-induced myofibroblast differentiation, a knockdown of Gα12 alone also had a moderate inhibitory effect, whereas knockdown of Gα13 had not (Fig. 4). It is noteworthy that while both Gα12 and Gα13 are linked to RhoA activation, they may recruit different Rho guanine exchange factors (GEFs) [22-25], and they may couple to different GPCRs [50], molecular mechanism of which has been under investigation [51]. Our studies have not revealed the role of Gαi and Gαq/11 in TGF-β-induced myofibroblast differentiation (Figs. 2, 3). This, however, does not negate their significance for fibroblast biology, given the established importance of Gαi/Gβγ and Gαq/11 signaling in cell proliferation, migration and contraction – all critical for the function of fibroblasts in wound healing and pathogenesis of tissue fibrosis. Finally, inhibition of TGF-β-induced myofibroblast differentiation by cholera toxin (Fig. 1) was intuitively expected given the reported inhibitory effects of cAMP-promoting agonists (i.e. prostaglandin E2, prostacyclin, adrenomedullin) [26-29]. However, to our knowledge, this is the first direct demonstration of regulation of TGF-β-induced myofibroblast differentiation by Gαs without affecting the proximal TGF-β signaling (Smad2/3 phosphorylation).

Our results suggest that a dependence of TGFBR1 and Smad3 expression on Gα12/13 (Fig. 5) could be one mechanism by which Gα12/13 control TGF-β-induced myofibroblast differentiation, which will be further evaluated in the future by forced overexpression of Smad2/3 and TGFBR1. However, other possibilities of regulation of Smad2/3 phosphorylation may exist, i.e. at the level of interaction of TGFBR1 with Smad2/3. For example, it was shown that small GTPase RhoB (but not RhoA) interacts with Smad3, blocks the interaction of Smad3 with TGFBR1and prohibits its phosphorylation [40]. The mechanism by which Gα12/13 control Smad2/3 signaling requires further investigation. Together, this study describes a novel crosstalk between TGF-β and G protein signaling in the context of myofibroblast differentiation and encourages new investigations on this crosstalk in other cellular functions of TGF-β.

## Acknowledgements

This study was supported by NHLBI Award R01 HL149993 (to NOD) and NINDS Award R01 NS111943 (to AAM).

## Notes

Funding: This study was supported by NHLBI Award R01 HL149993 (to NOD) and NINDS Award R01 NS111943 (to AAM).

### Competing Interest Statement

The authors have declared no competing interest.

